# CDR Conformation Aware Antibody Sequence Design with ConformAb

**DOI:** 10.1101/2025.11.12.688095

**Authors:** Imee Sinha, Samuel Stanton, Stephen Lillington, Sarah A. Robinson, Santrupti Nerli, Karina Zadorozhny, Joseph Kleinhenz, Homa MohammadiPeyhani, Michael Dillon, Yongmei Chen, Jack Bevers, Yan Wu, Andrew M. Watkins, Henri Dwyer, Richard Bonneau, Kyunghyun Cho, Franziska Seeger, Vladimir Gligorijević, Simon Kelow

**Affiliations:** Prescient Design, Genentech, South San Francisco, CA, USA; Department of Antibody Engineering, Genentech, South San Francisco, CA, USA; Center for Data Science, New York University, New York, NY, USA; Department of Computer Science, Courant Institute of Mathematical Sciences, New York University, New York, NY, USA; Canadian Institute for Advanced Research, Toronto, Ontario, Canada

## Abstract

Antibody lead optimization methods seek to enhance a lead candidate’s therapeutic properties through targeted sequence mutation. However, the mutations introduced during this process can inadvertently induce structural changes that disrupt binding, particularly by altering CDR loop conformations which destabilize CDR-target binding interactions. To address this, we present ConformAb, a guided discrete diffusion model for designing antibody sequences that explicitly conform to the CDR canonical conformation of the lead. With seed sequences as the starting point, we show that our method is capable of generating 3− 5× better binders that conform to the same CDR backbone structure in a one-shot design setup, and in some cases, binders with better affinity than seed repertoire picks without any target-specific data included in training. Across the targets tested, ConformAb demonstrated binding rates ranging from 15 −60% in wet lab experiments, a result obtained using fewer than 100 designs for each target. ConformAb offers a unique one-shot approach for antibody lead optimization in data-scarce scenarios where experimental/repertoire data cannot be leveraged for model training.

## 1 Introduction

Among therapeutic modalities, antibodies are highly effective biomolecules that selectively target a broad range of disease-associated proteins in cancers, autoimmune disorders, and infectious diseases [7, 32, 48]. Antibodies recognize and bind antigen proteins through six specialized loops (three originating from each antibody variable domain) called complementarity-determining regions (CDRs) which are evolved to effectively bind a wide variety of targets. [12]. Despite the vast variability in antibody sequences, the conformations of CDR backbones fall into a well-defined set of canonical clusters for most CDRs, enabling effective structure prediction from sequence data. Canonical clusters describe groups of CDR loops with the same length, nearly identical backbone structures, and shared sequence motifs [28, 35, 37]. These forms were first identified in antibody structures primarily from the research of Cyrus Chothia and Arthur Lesk [5, 9, 10], and spurred a systematic categorization that has been refined and expanded by other researchers [4, 35–39, 43, 49, 50]. In the most recent update in 2022, Kelow *et al* [28] provided a classification of antibody CDRs utilizing the DBSCAN algorithm and the maximum dihedral angle metric to provide 52 clusters, of which 16 were new and 36 remained the same as defined in North *et al* in 2011 [37], while singleton clusters and clusters with poorly resolved structures were discarded.

These structural insights are essential for antibody structure prediction and in lead optimization of antibody drug candidates using computational and deep learning methods [1,3,17,24,26, 31,41,47]. A key challenge in early-stage drug discovery is the frequent lack of data on a lead antibody candidate, such as antibody structural data, or associated repertoire data (i.e., naturally occurring antibody sequences selected from large-scale immune sequencing). This data scarcity limits current deep learning based antibody design frameworks that utilize antibody or antigen structure. For instance, structure-conditioned sequence design methods [13,25,34] are generally inapplicable as they require an initial structure of either the antibody or the antibody-antigen complex. Sequence-structure co-design methods [11,27,47] are difficult to apply successfully, because they face the hard problem of accurately predicting the correct antibody-antigen complex structure from sequence information alone to guide design. The alternative, protein language models (pLMs) [16, 19, 33, 42], can generate libraries of natural, expressible antibody sequences without a structural starting point while using the lead candidate sequence as the starting seed. However, this advantage comes at a cost, as pLMs often lack a structural prior to guide generation towards the paratope-epitope fit, producing sequences that are not always optimized for a specific binding mode [15, 45]. Consequently, sequences designed by pLMs may optimize for evolutionary fitness over CDR-epitope fitness and cause CDR loop conformational changes that disrupt target interactions.

To address this challenge, we present ConformAb, a fit-to-purpose model for predicting the canonical conformations of CDRs from the Fv sequence and for designing antibody sequences that explicitly preserve the CDR canonical conformation of the seed binder. In the absence of seed repertoire data or affinity labels for seed variants, our method offers a reliable solution for one-shot antibody design of sequences that are likely to retain both the conformation and function of the original antibody, increasing chances of binding and therapeutic effect.

ConformAb employs a guided discrete diffusion model [21] with masking noise to generate antibody sequences, jointly learning with six CDR-specific classifiers. During generation, the model is guided to produce sequences which conform to the seed canonical class probabilities for each CDR. With seed sequences known to bind three different antigen targets as the starting point, we show that ConformAb generated designs achieve binding rates ranging from 15-60% for all of the targets and demonstrate a 3−5× improvement in binding affinity over the seed for two of the three targets. Experimental structure prediction done for binders of two targets further reveal that ConformAb designs maintain the backbone CDR canonical conformation of the seed, even in the presence of biophysically non-trivial mutations. We also demonstrate the utility of ConformAb as a scoring function: on an internal dataset, ConformAb successfully predicted binders with an accuracy of 68% for designed sequences. Overall, our results highlight ConformAb’s potential as a naive binder generation method to accelerate antibody lead optimization and hit expansion.

## 2 Results

### 2.1 Folding methods recapture native antibody CDR canonical conformations

We trained on two datasets, the SabDab [14] set of sequences and the pOAS [29, 40]. While SabDab represents the collection of sequences from PDB crystallographic and cryo-EM data, pOAS is the set of publicly released sequences obtained from repertoire sequencing studies, without accompanying crystallographic structural data. Recent work has shown that ABB2 [2] predicted structures of the pOAS sequences majorly correspond to the existing CDR canonical cluster structural space [20]. In addition, we validated that ABB2 folded structures accurately recapture native canonical conformations for both the SabDab dataset as well as the ABB2 validation set. Table 1 shows the accuracies of recovering native CDR canonical conformations for sequences folded with ABB2 for the SabDab dataset and the ABB2 validation set. Accuracy is based on the dihedral clustering metric used in Kelow *et al* [28], with a threshold of 35 degrees as a cutoff for recovery of the native CDR conformation. Accuracies show that folding models reliably recapture native CDR canonical conformations, which is key to training the ConformAb model using canonical cluster labels from both the SabDab crystallographic dataset and the folded pOAS dataset, allowing access to a large amount of sequence diversity in training.

**Table 1.** Accuracy of recovering native CDR canonical conformation from ABB2 folded structures. Across all CDRs, accuracies are high, the lowest being for H3.

|  | **H1** | **H2** | **H3** | **L1** | **L2** | **L3** |
| --- | --- | --- | --- | --- | --- | --- |
| **ABB2 validation set** | 0.93 | 0.83 | 0.73 | 1.00 | 0.83 | 0.87 |
| **SabDab dataset** | 0.89 | 0.90 | 0.86 | 0.89 | 0.87 | 0.83 |

### 2.2 ConformAb trained on the joint PDB and pOAS datasets provides high accuracies for predicting antibody CDR canonical conformations

Figure 1 shows the canonical cluster landscape for SabDab (PDB) and the folded pOAS dataset. Canonical clusters (x-axis) are displayed beginning with the CDR identifier (e.g., H1), followed by the sequence length (e.g., 13), and finally the specific cluster within that length-for example, H1-13-1. Cluster names match those established in Kelow *et al* [28]. Both the PDB and pOAS datasets contain defined clusters (or classes) for all CDRs, as well as some undefined “noise” clusters (denoted by *). CDR H3, in particular, has the largest proportion of sequences within these noise clusters. Unlike defined clusters, noise clusters do not correspond to a consistent structural pattern, instead, they contain CDR sequences that do not fit into any of the defined structural clusters for that length. Overall, PDB and pOAS canonical cluster landscapes exhibit similar trends, however, the pOAS consistently shows a higher proportion of sequences assigned to the noise cluster for H3, particularly for longer CDR lengths. We used the PDB dataset and the joint PDB+pOAS dataset to train two independent models to predict canonical conformations of the six CDRs of an antibody sequence.

**Figure 1.**
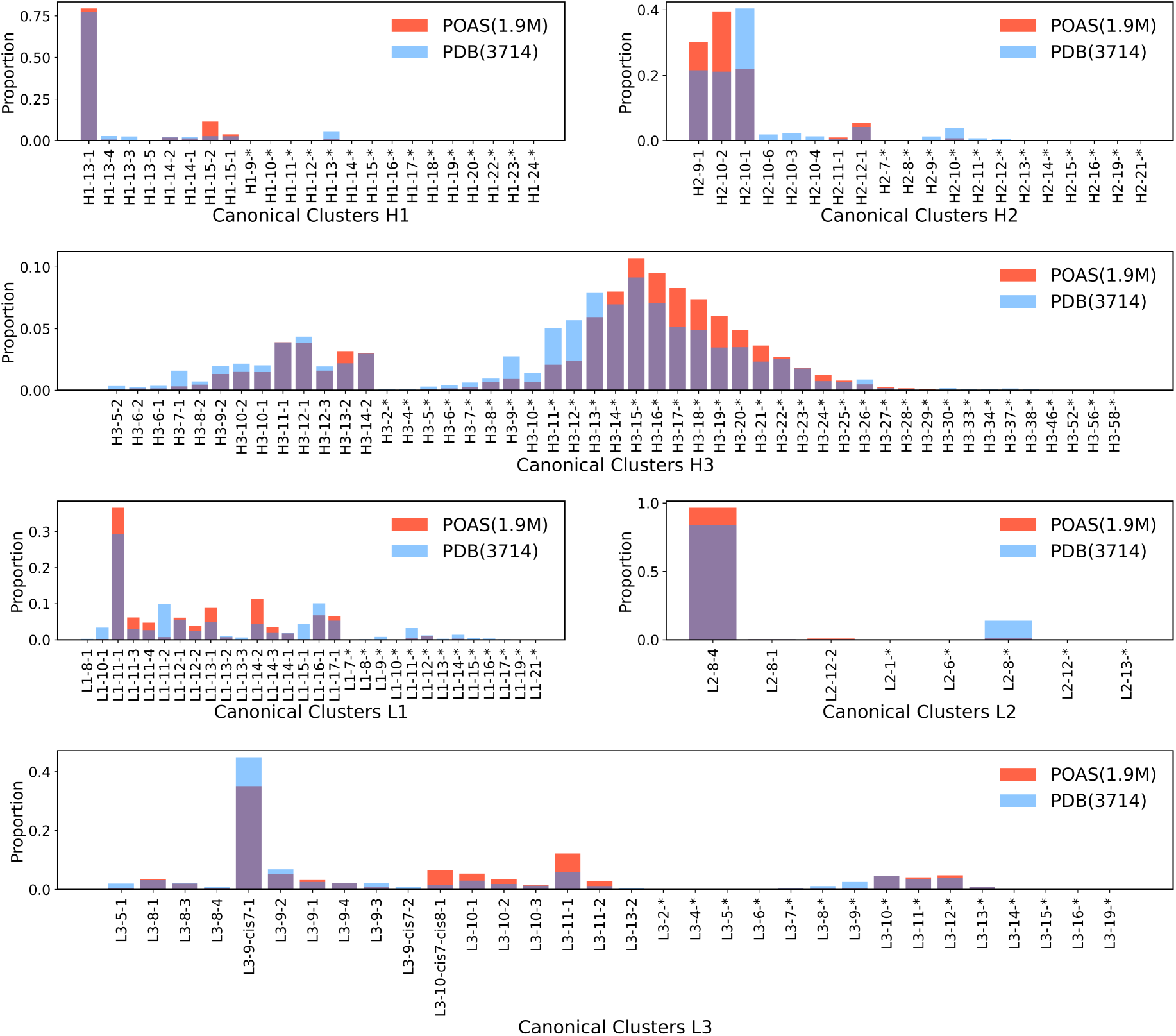
Canonical cluster landscapes for SabDab and folded pOAS structures. Panels show the proportion of the total dataset assigned to each cluster (y-axis) for the six CDRs. Canonical clusters are shown on the x-axis, labeled by the CDR type (e.g., H1), sequence length (e.g., 13), and the cluster ID within that length-for instance, H1-13-1. Data represents 1.9 million folded pOAS sequences and 3,714 structures from the SabDab dataset. Clusters ending in “*” denote noise clusters.

In Table 2, we show the test accuracies of the ConformAb model trained on (i) the PDB dataset and on (ii) the PDB+pOAS dataset. The performance of the model critically improves for H3 and L3 CDRs with the addition of data from the pOAS dataset, notwithstanding the increase in the number of canonical clusters within the pOAS dataset (Table 5). However, for both models, the accuracy of H3 and L3 predictions remain significantly lower compared to other CDRs, likely due to the greater prevalence of noise clusters in H3 and L3, making prediction more challenging. Table 5 provides the total number of examples present for training the PDB model and the PDB+pOAS model as well as the total number of clusters and noise clusters for each CDR within both datasets. The increase in the number of noise clusters in the pOAS data is reflected in the slight decrease of accuracy for H1 for the PDB+pOAS model on the PDB test set (the total number of clusters increase from 14 to 22 for H1 by addition of 7 noise clusters). Both the PDB and PDB+pOAS models perform better on test sets drawn from their own respective datasets. Notably, the PDB+pOAS model achieves comparable (or higher) performance to the PDB model across both test sets, and significantly outperforms it on H3 and L3 predictions for both test sets.

**Table 2.** Accuracies of PDB and PDB+pOAS model for canonical cluster prediction. Two types of test sets were used: one derived solely from the PDB dataset and the other derived solely from the pOAS dataset. Notation such as 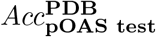 indicates the performance of the PDB-trained model on the pOAS test set. The last two columns report the accuracy of the PDB+pOAS model, which is comparable or higher than the PDB model across both test sets.

| CDR | $Acc_{\text{PDB test}}^{\text{PDB}}$ | $Acc_{\text{pOAS test}}^{\text{PDB}}$ | $Acc_{\text{PDB test}}^{\text{PDB+pOAS}}$ | $Acc_{\text{pOAS test}}^{\text{PDB+pOAS}}$ |
| --- | --- | --- | --- | --- |
| **H1** | 0.78 | 0.69 | 0.76 | 0.89 |
| **H2** | 0.81 | 0.59 | 0.83 | 0.96 |
| **H3** | 0.46 | 0.16 | 0.74 | 0.81 |
| **L1** | 0.76 | 0.72 | 0.82 | 0.95 |
| **L2** | 0.83 | 0.74 | 0.80 | 0.96 |
| **L3** | 0.63 | 0.27 | 0.72 | 0.85 |

### 2.3 ConformAb predicted canonical conformations compare favorably with ESMFold and ABB2 folded structural assignments

To investigate potential biases arising due to the choice of folding method, we compared whether the ConformAb model - trained using sequences and canonical class labels from ABB2 folded structures - exhibited an over-representation of certain canonical clusters when compared to labels derived from a different folding method.

To evaluate this, we benchmarked the structurally annotated canonical classes from ABB2 [2] as well as ESMFold [30] folded structures to ConformAb based predictions for a randomized test set of 2000 antibody sequences from pOAS (held out from ConformAb training). Canonical classes of the ABB2 and ESMFold folded structures were determined using the dihedral angle based clustering method 4.1. We compared the frequency of antibody CDRs assigned to each canonical class by the three methods (Figure 2a) and, given the low frequency of some class predictions, we also represented this data as a percentage to enable better visualization of differences within low-represented classes (Figure 2b).

**Figure 2.**
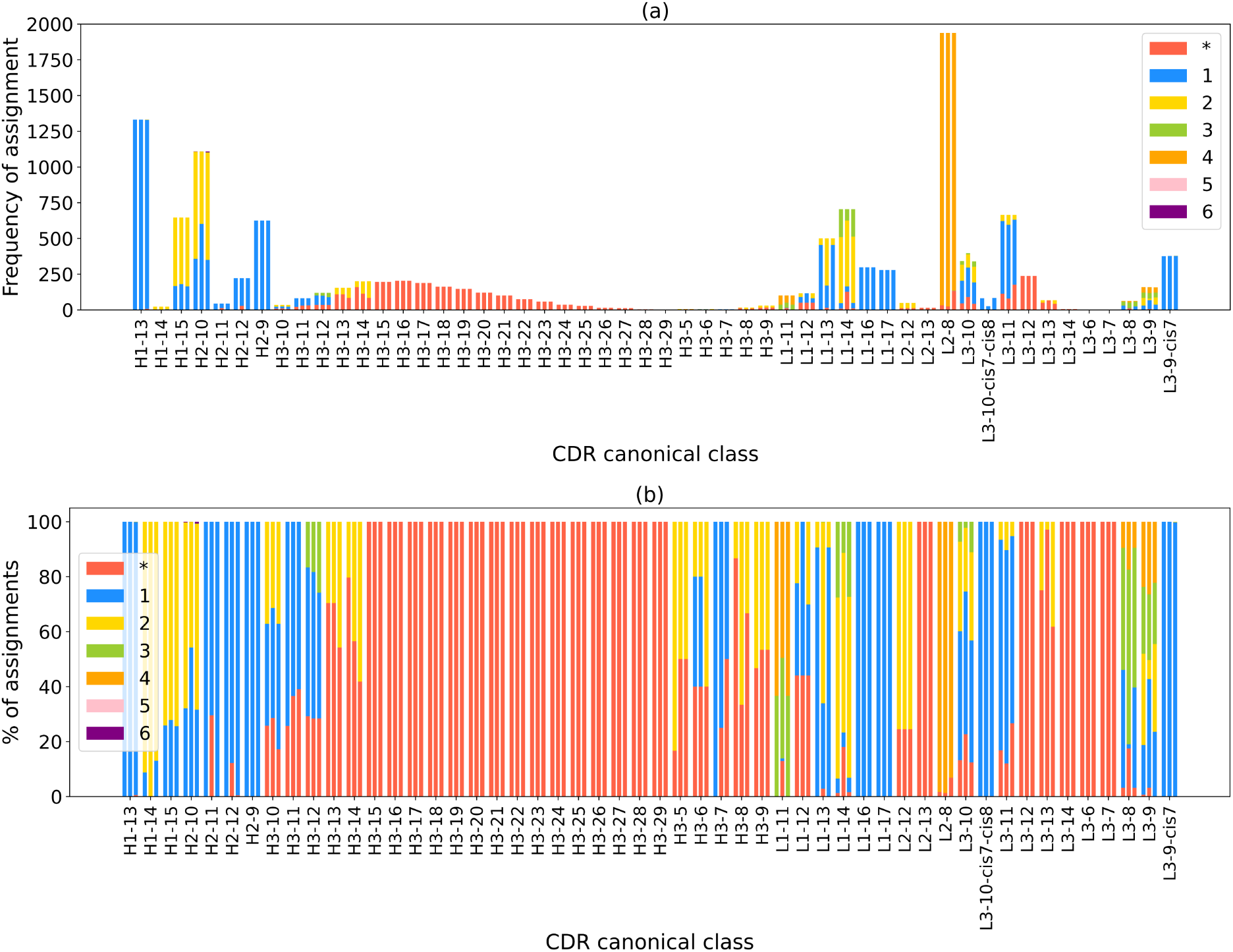
Comparison of ConformAb with ESMFold and ABB2 structural assignments. Subplot (a) shows the frequency of assignment of CDR sequences to a cluster, out of 2000 total sequences. Each color represents a distinct cluster I.D. within a given CDR length (depicted on X axis). Three bars within each CDR length represents the assignment from ABB2, ESMFold and ConformAb respectively from left to right. Subplot (b) shows the percentage allocation of clusters across CDR lengths to highlight variations within underrepresented classes.

Overall we see fairly high levels of agreement between the methods. Many CDR lengths contained only one canonical class prediction by all three approaches; for example, H2-9 (all H2-9-2), L1-16 (all L1-16-1) and L1-17 (all L1-17-1). All CDR H3 longer than 14 amino acids were assigned entirely to noise clusters as no canonical classes exist for H3 with length*>*14 within this dataset, as is the case for light chain CDRs L2-13, L3-6, L3-7, L3-12 and L3-14. L3-9 had the most variation in canonical conformations, with all four canonical classes predicted by each method, in addition to noise clusters. Overall, ConformAb predictions aligned closely with ABB2 structural labels over the set of 2000 sequences, especially for H1, H2, L1, and L2 (≥ 95% agreement), while H3 and L3 showed slightly lower agreement between ConformAb and ABB2 labels (86% and 90% agreement, respectively). ConformAb showed slightly lower agreement with ESMFold structural labels, with drops in percentage agreement for H2 (84%), H3 (87%), L1 (75%), and L3 (78%), while H1 (93%), and L2 (94%), showed strong agreement between labels.

To highlight differences within rare clusters that appear in limited number of sequences, we analyzed the percentage allocation across clusters (Figure 2b) which improved visibility of variations within underrepresented classes. Agreement between ConformAb and ABB2 for these classes was generally expected given ConformAb was trained on ABB2 folded structures. We observed some differences primarily across CDR H3, with varying levels of noise cluster assignments. This was particularly evident for H3-7 where 100% of ABB2 structures were classified as H3-7-1 but 50% were predicted as noise by ConformAb (Figure 2b). In contrast, differences between ABB2(or ConformAb) and ESMFold were notable for lengths H1-14, H2-10, H2-11, H2-12, L1-13 and L1-14. For H1-14, 100% of ESMFold models adopt canonical class H1-14-2 rather than H1-14-1 which was adopted by ConformAb and ABB2. For H2-11 and H2-12 CDRs, only ESMFold structures were classed into noise clusters. Across the light chain CDRs, ESMFold structures were more frequently assigned to noise clusters than the other two methods (e.g. L1-11, L1-14, L3-8, L3-10, L3-13). Interestingly, ESMFold structures are less likely to adopt L3-10-cis7-cis8 canonical class but instead favor other L3-10 canonical forms. It’s worth noting that many of these clusters showing large differences across the three methods are observed within a smaller set of sequences — often fewer than 10 — and are likely underrepresented within the pOAS data.

### 2.4 ConformAb produces diverse sequences while preserving canonical conformation

ConformAb designs antibody CDR sequences using a seed binder as the starting point. The binding of the initial seed ensures that a functional binding mode exists, and designed sequences are generated to maintain the original CDR conformation while introducing diversity via mutations. When the seed contains canonical CDRs, mutations can be applied to all CDR regions to introduce diversity. In contrast, for seeds with non-canonical CDRs (CDRs assigned to noise clusters), mutations are selectively restricted to canonical CDR regions, leaving non-canonical CDR sections intact to preserve the “noisy” binding mode of the original seed.

To illustrate, antibody sequences were generated for 4 seeds with canonical CDR conformations. Each of the six CDRs was masked (in independent and pairwise combinations), and sequences were generated to introduce diversity within masked regions. The resulting edit distances of CDR designs, shown in Figure 3c (average edit distances reported in Table 3), depends on the CDR sequence and its canonical cluster. While the edit distances in Figure 3c (and Table 3) relate to individual CDRs, it indicates that mutations spanning multiple CDRs generate substantial diversity, often pushing total edit distances to well beyond 25-30. During ConformAb generation, these edits can be limited to select CDR regions, or distributed widely across all CDR regions. In this work, we have not evaluated framework mutations based designs, although those designs are possible within the ConformAb framework.

**Table 3.** Average edit distances observed across designs. Average edit distances of CDR sequences generated for seeds 1a7, 1dqm, 1jfq, and 3v0w are shown for each CDR. For each seed, the canonical cluster of the corresponding seed CDR is listed directly below the edit distances.

| PDB | H1 | H2 | H3 | L1 | L2 | L3 |
| --- | --- | --- | --- | --- | --- | --- |
| 1a7p | 7.14 | 5.6 | 6.24 | 3.95 | 5.78 | 5.85 |
|  | (H1-13-1) | (H2-9-1) | (H3-10-1) | (L1-11-2) | (L2-8-4) | (L3-9-4) |
| 1dqm | 6.37 | 3.02 | 5.43 | 3.44 | 5.94 | 5.02 |
|  | (H1-13-1) | (H2-9-1) | (H3-7-1) | (L1-11-1) | (L2-8-4) | (L3-9-cis7-1) |
| 1jfq | 5.88 | 8.27 | 7.45 | 5.27 | 4.48 | 4.98 |
|  | (H1-13-1) | (H2-10-1) | (H3-14-2) | (L1-11-2) | (L2-8-4) | (L3-9-cis7-1) |
| 3v0w | 4.09 | 8.19 | 7.83 | 6.70 | 3.61 | 4.86 |
|  | (H1-13-1) | (H2-12-1) | (H3-11-1) | (L1-11-1) | (L2-8-4) | (L3-9-cis7-1) |

**Figure 3.**
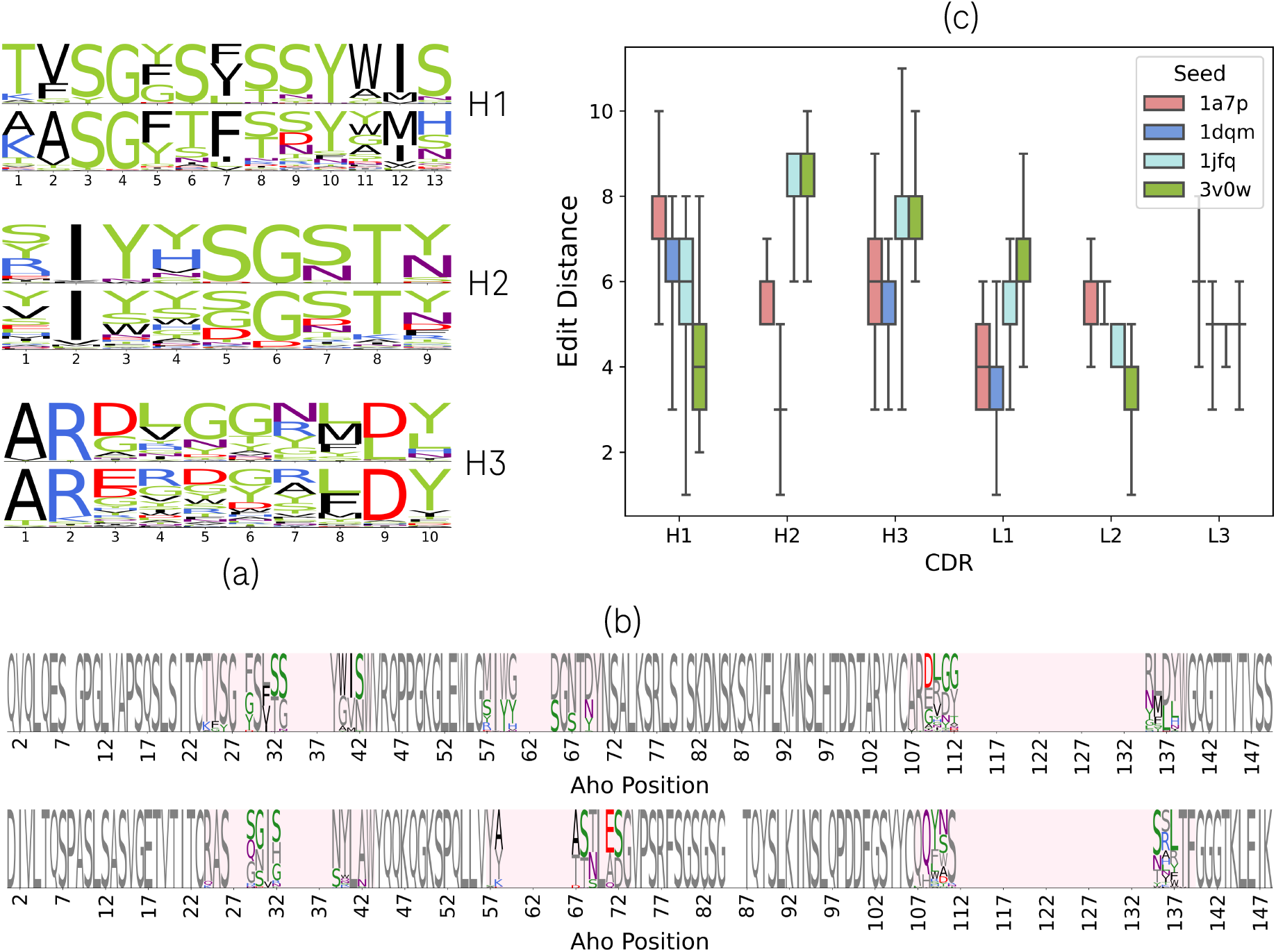
Sequence generation with ConformAb. Subplot (a) presents a comparison of CDR motifs for ConformAb generated 1a7p designs (top) and their corresponding canonical cluster motifs (bottom) across the CDRs H1, H2, and H3. Residue colors follow scheme described in Kelow *et al*. [28]. In (b), sequence logos display the design variability across CDR regions of both heavy(top) and light(bottom) chains for 1a7p designs, with mutations shown in color and the seed sequence in gray. Mutations are restricted to the CDRs regions highlighted in lavender. Subplot (c) summarizes distribution of edit distances per CDR for all seeds using box plots. Each box plot displays the median (center line), interquartile range (box), and the full data range from minimum to maximum (whiskers). The highest diversity is observed in CDR H3, while diversity in other CDRs are influenced by their respective seed CDR sequences and canonical cluster.

Figure 3a and 3b illustrates the CDR and sequence logo plots respectively for sequences generated from starting seed 1a7p. The generated CDR sequences in Figure 3a (top) strongly resemble their canonical cluster motif (bottom) [28], with diversity being constrained by the patterns present in starting seed CDR sequence. For instance, although both 1a7p and 3v0w belong to H1-13-1 canonical cluster, 1a7p exhibits greater mutational diversity on CDR H1 and an higher average edit distance (7.14) compared to 3v0w (4.09) (Table 3).

Some ConformAb designs feature “locked-in” effect, where a single residue, or groups of residues, dominate, repeating themselves through all designs. This is expected, as ConformAb inherits dominant patterns from the canonical cluster, however this also reduces diversity of the generated designs. To minimize locked-in effect, masking should be strategically applied, particularly at positions that contain dominating motifs. Random sampling (all CDR positions masked), on the other hand, is more likely to produce designs that consistently contain repeating “locked-in” patterns in every sequence.

Figure 3b illustrates the sequence logo for 1a7p designs created via ConformAb. On H1, dominant “locked-in” mutations included G to W at Aho position 40 (corresponding to position 11 on the CDR logo plot), G to S at Aho position 33 (position 9 on the CDR logo plot), V to I or M at Aho position 41 (position 12 on the CDR logo plot), and N to S at Aho position 42 (position 13 on the CDR logo plot). These mutations follow position-specific residue probabilities for H1-13-1 canonical cluster [28], hence the tendency of designs to mutate to these residues.

### 2.5 ConformAb serves as ranking model to score designs and successfully performs naive binder generation

To assess ConformAb’s potential as a scoring method, we tested it on a set of designs generated for EGFR, IL6, and OSM targets via lab in the loop (LITL) antibody discovery experiments described in Frey *et al* [18]. In total, 1609 designs (consisting of 958 binders and 651 non-binders/non-expressors) were generated and experimentally tested for binding to the above targets, using ML methods and datasets detailed in Frey *et al* [18] (no ConformAb generated designs were included in this analysis). Figure 4a visualizes ConformAb output KL divergences of seeds to designs for these sequences. Here, the KL divergence quantifies the difference between ConformAb predicted canonical class probabilities of each design from its seed. A divergence close to zero suggests strong alignment of seed and design class probabilities, indicating similar CDR backbone conformations. As seen in Figure 4a, binders exhibit a pronounced peak at zero KL divergence, whereas non-binders display a broader KL divergence distribution with lower and wider peaks, indicating significant differences from the seed. Across the six CDRs, the differences in peaks are most pronounced in L1, L3, and H1, indicating that binders show prominent variations from non-binders in these regions, while L2 exhibits the least difference amongst binders and non binders. For these set of designs, we computed the ConformAb binding score by comparing if their ConformAb predicted canonical class matched that of the seed. Designs that matched were labeled ‘predicted binders,’ while those that did not were labeled ‘predicted non-binders’. Table 4 shows contingency table of predicted vs. true binders (and non binders) for this experiment. ConformAb achieved 68% accuracy in predicting binders using this approach, with 0.72 precision and 0.76 recall, meaning it identified positives pretty accurately, and identified most of the true binder designs. Further, a chi-squared test of independence showed a significant association between ConformAb predictions and experimental binding outcomes (*χ*^2^ = 170.23, *p <* 0.05), indicating strong agreement.

**Table 4.** ConformAb Binder Prediction Performance. Contingency table showing ConformAb predicted binders and non-binders vs. true binders and non-binders from the LITL set of generated designs.

|  | Predicted Non-binder | Predicted Binder |
| --- | --- | --- |
| Non-binder (true) | 363 | 288 |
| Binder (true) | 227 | 731 |

**Figure 4.**
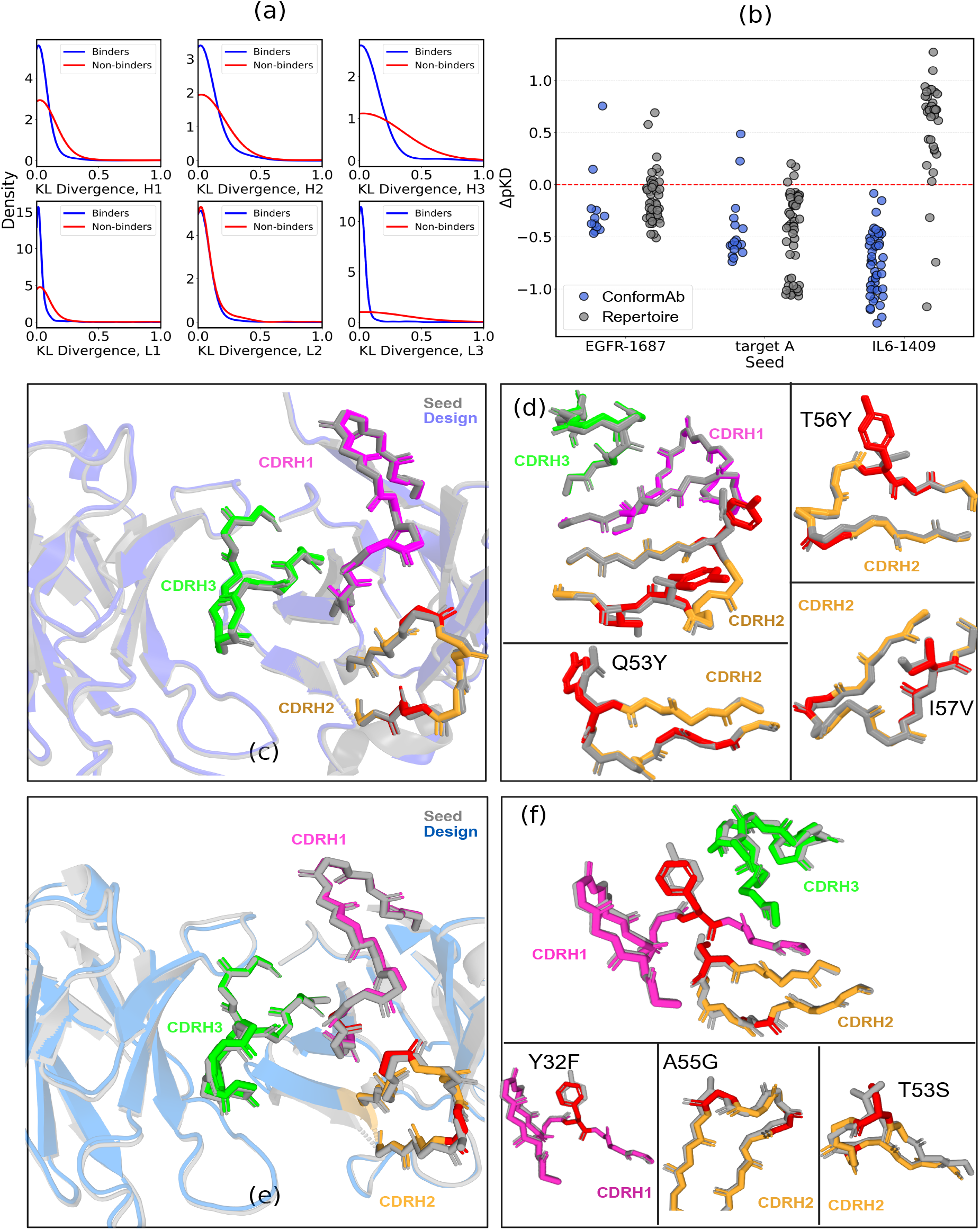
Naive binder generation and scoring. (a) KL divergence of LITL designs from seeds, showing binders are more concentrated at zero than non-binders. (b) Experimental Δ*pK*_*D*_ for ConformAb designs and repertoire picks targeting EGFR, target A, and IL6; the red dashed line (y=0) marks the seed baseline. (c, e) Structural overlays of seed (gray) and design (blue) for EGFR and IL6, respectively. (d, f) Key mutations (red) for the EGFR (Q53Y, T56Y, I57V) and IL6 (Y32F, A55G, T53S) binder designs. For all designs, CDRs H1-H3 are in magenta, orange, and green while mutations are in red. CDR backbone is rendered as sticks while side chains are hidden, except to highlight mutations in panels d,f.

In a separate experiment, we evaluated ConformAb’s performance in naive binder generation task by generating designs for three targets: IL6, EGFR, and target A (an internal target, anonymized here), based on lead antibody sequences as seed. For IL6 and EGFR, a single seed sequence (IL6-1409 and EGFR-1687 respectively) was tested in each case, whereas target A was tested using two distinct seeds. Figure 4b shows results for binder generation and corresponding SPR based binding affinity of designs. Here, binding affinity is defined as the negative log-transformed equilibrium dissociation constant: 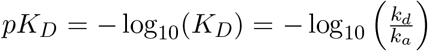, with *kd* and *ka* representing the dissociation and association rate constants, respectively. Each design set includes antibody designs derived from an antibody ‘lead’, a known antigen binder with given *pK*_*D*_, and binding affinity differences, Δ*pK*_*D*_, are computed relative to this lead (also termed ‘seed’). For the three antigen targets, ConformAb successfully generated binders on the first generation without any auxiliary target-specific data. In the case of EGFR and target A, ConformAb produced two binders having significant improvement in binding affinity over the seed during the first round of generation. Success for target A was notably seed-dependent, as the second seed for target A failed to produce any binders. The overall binding rate across all targets was 22%; designs based on the successful seed for target A yielded a 19% binding rate, while IL6 and EGFR yielded 57% and 15% binding rates, respectively. Importantly, none of these designs required any data beyond the seed sequence for binder generation. In the absence of random mutagenesis data as the ideal baseline, we used repertoire-based picks for comparative benchmark of Δ*pK*_*D*_. Repertoire picks are sequences from the seed clonotype that are presumed to bind the target antigen and were selected using internal scoring methods based on trained protein language models. Repertoire picks were sampled across a range of scores to reduce potential bias or inaccuracies in the scoring functions. Notably, the top ConformAb binder outperformed all the repertoire picks for EGFR and had *>* 5× improvement in binding affinity over seed, (Δ*pK*_*D*_ = 0.75) (Figure 4b). For target A, the top ConformAb binder had a *>* 3× improvement in binding affinity over seed (Δ*pK*_*D*_ = 0.48), and was ~ 2× better than the top repertoire picked binder sequence. For IL6, ConformAb achieved a high binding rate but did not surpass the seed or any of the repertoire picks in affinity. These findings demonstrate that ConformAb can generate binders that are competitive with, and sometimes superior to, repertoire informed picks without any auxiliary data except the seed sequence.

To examine underlying backbone conformations, we determined the crystal structures of the original lead and the top performing ConformAb binders for both EGFR and IL6. For EGFR, the seed and binder crystal structures were determined to 1.78 Å and 2.4 Å resolution respectively. The optimized EGFR binder incorporates three mutations in the CDR H2 loop - Q53Y, T56Y, and I57V, designed to promote binding. Figures 4c-d shows the EGFR seed and design structures overlaid, and the CDR backbone structures which conform to each other. CDR backbones are rendered as sticks to show alignment, with side chains hidden for clarity-except for the mutations highlighted in Figure 4d. Figure 4d shows each of the three mutations in detail, showing the side chain of the original residue as well as mutated residue (all other side chains hidden for clarity). Despite the introduction of two biochemically disparate and large, ring-based mutations (Q53Y and T56Y), the CDR backbone conformation of the design remained unaltered relative to the original seed. While the backbone is conserved, the side chain mutations introduce new hydrophobic interactions. For IL6, crystal structures for the lead IL6 Fab as well as the top IL6 binder were determined to a 1.9 Å and 2.61 Å resolution respectively for structural comparison. Three mutations - Y32F, A55G, and T53S, were introduced in CDR H1 and H2 of the heavy chain to promote binding. Figure 4e and Figure 4f shows the seed and the design structures overlaid, their CDR backbones in stick representation, and the mutations in red. Side chains are hidden for clarity, except in Figure 4f, where they are shown only for the mutated residues and their original counterparts. Throughout the design, and across all CDRs, the backbone conformation remains unaltered relative to the seed, demonstrating ConformAb’s ability for CDR backbone conforming binder design.

## 3 Discussion

Designing antibody sequences based on a lead candidate, especially in low-data regimes, remains a significant challenge in therapeutic antibody development. Because the conformation of CDR loops affects how antibodies bind, optimizing antibody sequence without explicitly preserving structural conformation can risk altering these loops and reduce binding.

To address this, we introduce a structure-aware sequence design framework that explicitly conditions designs on the CDR loop conformations of the seed binder. Our results demonstrate that ConformAb enables a range of sequence edits while conforming to seed CDR canonical structure via generation. Using ConformAb, mutations can be focused within a single CDR or distributed across multiple CDRs, resulting in total edit distances of 25–30 when all six CDR loops are mutated. Amongst them, CDR H3 tends to show the most variation during sequence generation. Further, designs across all CDRs closely mirror their respective cluster motifs, as well as the original seed CDR sequence.

Because it encodes a learned structural prior through canonical conformations, ConformAb provides a method to rank designs that are structurally similar to seed. Using KL divergence between ConformAb class probabilities of seeds and designs, we found that binders have lower divergence from the seed than non-binders. During naive binder generation, ConformAb successfully generated binder sequences for three antigen targets tested in the first round of design using only the seed sequence, without any supporting repertoire data. Notably, for both EGFR and target A, ConformAb produced binders with significantly improved binding affinities compared to their respective seeds. Structural comparison of the top EGFR binder with respect to the seed shows that while the binder introduces novel, non-conservative mutations, it adheres to the seed backbone structure at the same time.

Overall, ConformAb presents a promising approach for antibody sequence design in data-limited scenarios. Investigating how dynamics effect canonical class assignment and how canonical conformations correlate with downstream tasks such as epitope recognition and developability is a key next step. Expanding the model to incorporate emerging new canonical clusters remains critical to improve its accuracy. Additionally, integrating ConformAb with complementary predictive frameworks, such as those for developability and immunogenicity could unlock new strategies for multi-objective optimization in antibody design.

## 4 Methods

### 4.1 Canonical cluster dataset curation

In order to train the ConformAb model, we required sequences labeled with antibody CDR canonical conformations. Previous work by Kelow et. al [28] curated a dataset based on canonical conformations for the SabDab dataset [14]. This work derived the latest set of canonical conformations by clustering CDR dihedral features for all antibody structures available in the SabDab datset. To curate our dataset, we leveraged the cluster centers listed in the work by Kelow et. al to assign CDR structures to their nearest cluster center using a distance cutoff of 35 degrees. If a CDR structure is outside of the 35 degree cutoff, it is listed as noise. We do this assignment for each structure in SabDab and the folded pOAS [29, 40] to generate the data consisting of light and heavy Fv sequences and canonical cluster labels for each of their CDRs.

### 4.2 ConformAb model dataset curation

Table 5 summarizes the composition of the PDB and pOAS datasets used for model development across the 6 CDRs. The PDB+pOAS dataset contains a greater number of clusters than the PDB, due to additional noise clusters in the pOAS. To ensure balanced contribution from the PDB dataset, only a portion of the pOAS data was utilized during PDB+pOAS model development.

**Table 5.** Dataset Description. Columns list, for each CDR (column 1), the number of sequences used for the ConformAb model (column 2), total clusters (column 3), and noise clusters (column 4) in both the PDB dataset (columns 2-4) and the PDB+pOAS (columns 5–7) dataset respectively. The only increase in clusters within the PDB+pOAS dataset comes due to addition of noise clusters from the pOAS dataset.

| CDR | $N^{\text{PDB}}$<br>Dataset | $N^{\text{PDB}}$<br>clusters | $N^{\text{PDB}}$<br>noise clusters | $N^{\text{PDB+pOAS}}$<br>Dataset | $N^{\text{PDB+pOAS}}$<br>clusters | $N^{\text{PDB+pOAS}}$<br>noise clusters |
| --- | --- | --- | --- | --- | --- | --- |
| H1 | 3644 | 14 | 6 | 33111 | 22 | 14 |
| H2 | 3655 | 14 | 6 | 40404 | 20 | 12 |
| H3 | 3646 | 40 | 27 | 85373 | 44 | 31 |
| L1 | 3656 | 27 | 10 | 48821 | 30 | 13 |
| L2 | 3660 | 7 | 4 | 35277 | 7 | 4 |
| L3 | 3654 | 29 | 12 | 84172 | 31 | 14 |

In preparing the PDB dataset, we retained all structures except seed PDBs used for sequence generation (Section 2.4). Additionally, duplicate sequences across the dataset were eliminated, ensuring only unique entries were retained. Clusters containing fewer than four members were excluded from further analysis. For each cluster, we applied a randomized data-splitting approach with a 70:15:15 ratio for training, testing, and validation. These subsets were then merged across all clusters to construct the final dataset for model development and evaluation. This dataset was used to exclusively train the PDB model.

For the pOAS dataset, we selectively retained non VHH antibody sequences from human, mouse, rat, and rabbit species. For each CDR, only sequences with CDR lengths matching those in the PDB dataset were included, and duplicates were eliminated. Additionally, a holdout set of 2,000 sequences was randomly selected for folding method evaluation (section 2.3) Clusters containing fewer than four members were excluded from analysis. To ensure balanced representation across clusters, we applied the following downsampling schema.

**Clusters with greater than 100,000 members**: Retained 1% and split into training (60%), validation (20%), and test (20%) sets.

**Clusters with 10,000–100,000 members**: Retained 5%, and split into training (60%), validation (20%), and test (20%) sets.

**Clusters with 1,000–10,000 members**: Retained 80% and split into training (60%), validation (20%), and test (20%) sets.

**Clusters with 10–1,000 members**: Retained all the data, and split into training (80%), validation (10%), and test (10%) sets.

**Clusters with 4-10 members**: Retained all the data and split approximately into training (33.33%), validation (33.33%), and test (33.33%) sets.

The pOAS dataset curated above was subsequently merged with the PDB dataset across train, and validation sets for the training of the PDB+pOAS model. This downsampling strategy also ensured a balanced ratio of data from PDB crystallographic structures and pOAS sequences, ensuring a minimum limit on the PDB-to-pOAS data ratio while maintaining balanced cluster representation within the data.

The PDB model was trained solely on the PDB dataset, while the PDB+pOAS model utilized data from both the PDB and pOAS datasets. For the PDB+pOAS model, the train and validation splits from the PDB and pOAS data sources were concatenated separately for each split, ensuring no mixing between splits. Test sets from the PDB and pOAS were kept separate and independent to evaluate the performance of both models across two datasets. As a result, the accuracy of each model could be precisely evaluated on both PDB and POAS test sets respectively. Both models were trained for up to 1M steps, and the accuracy for each of the six CDRs was assessed.

### 4.2 ConformAb Model Architecture, Training, and Usage

The ConformAb model generates antibody sequences that preserve the CDR canonical conformations of a seed antibody by integrating discrete diffusion with classifier-guided sampling.

#### 4.3.1 Background: Discrete Diffusion for Antibody Design

Discrete diffusion models generate sequences by learning to reverse a masking process, where amino acids are gradually replaced with [MASK] tokens during training and then restored during generation [6, 23]. We apply classifier guidance using the diffusio**N O**ptimized **S**ampling (NOS) algorithm [21] to steer generation toward sequences preserving CDR canonical conformations. During each denoising step, NOS modifies token predictions using an explicit value function—in our case, the KL divergence between predicted CDR canonical class distributions of the design and seed. The NOS remasking approach also connects to recent advances in consistency models [44] and restart sampling [51], which similarly alternate between predicting a “clean” example from a corrupted example and re-corrupting the clean example for additional refinement.

#### 4.3.2 ConformAb Multi-Task Architecture

ConformAb combines sequence generation with CDR canonical class prediction through a shared encoder architecture (Figure 5). A key technique is joint training of diffusion and classification tasks with ensemble prediction heads for robust canonical class estimation [21].

**Figure 5.**
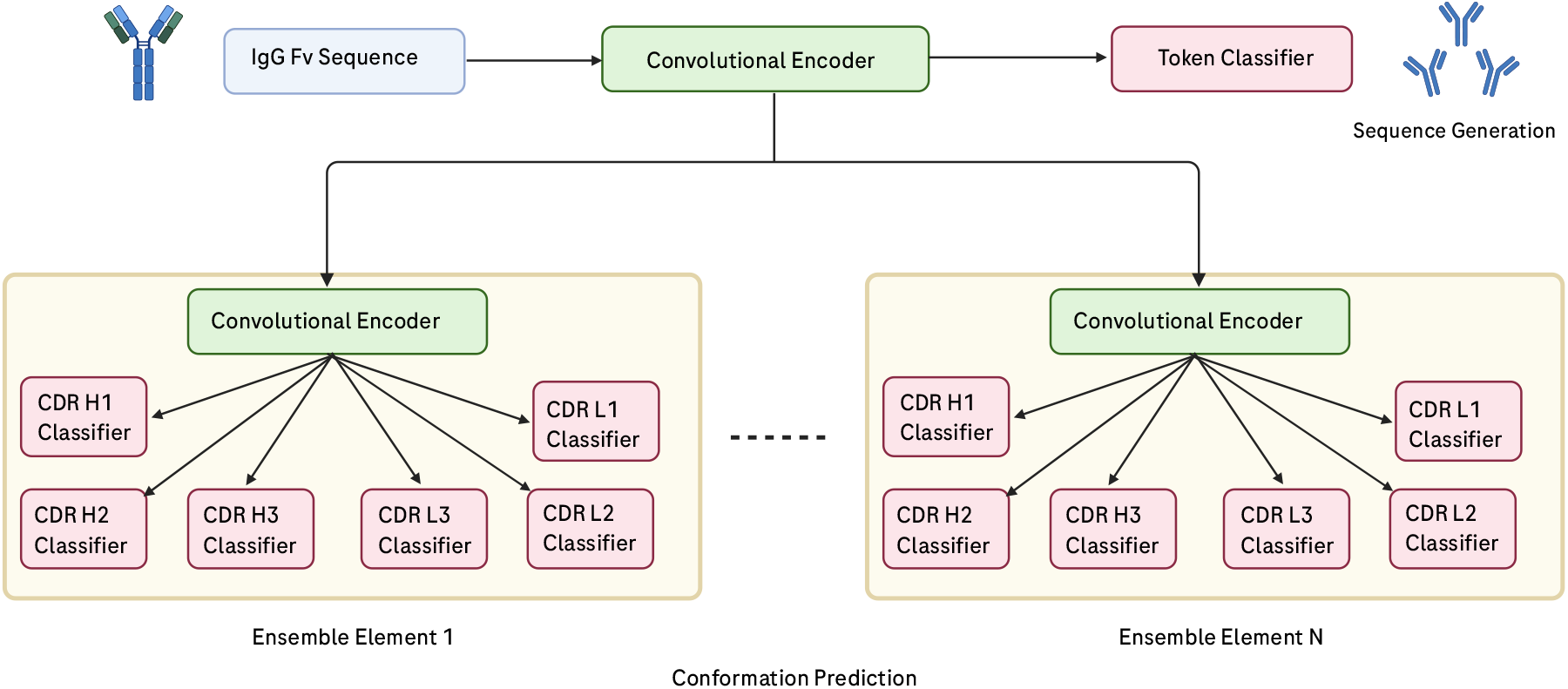
ConformAb Model Architecture. An input Fv (variable domain) sequence is processed by a shared convolutional encoder that feeds both the token prediction head for diffusion and ensemble classifier elements. Each ensemble element contains a branch-specific encoder and six CDR canonical class classifier heads. The classifier predictions provide value guidance during generation.

The architecture comprises three main components: (1) A **shared convolutional encoder** that processes input antibody Fv sequences and learns representations for both generation and classification tasks; (2) A **token classifier head** that performs the discrete diffusion denoising task, outputting amino acid logits for each sequence position; (3) An **ensemble of CDR classifier modules**, where each module contains a branch-specific encoder and six classifier heads predicting canonical classes for CDRs H1, H2, H3, L1, L2, L3. The ensemble design improves prediction robustness and provides epistemic uncertainty estimates for guidance under imperfect information. During generation, classifier predictions from all ensemble members inform the KL divergence guidance signal that steers the diffusion process toward preserving seed CDR canonical conformations.

#### 4.3.3 Training Techniques

**Round-Robin Partition Rebalancing** Standard balanced sampling approaches inadequately handle the extreme class imbalance in CDR canonical class datasets, where rare canonical classes can be underrepresented by orders of magnitude. We introduce **Round-Robin Partition Rebalancing (RPR)** (Algorithm 1), a stateful minibatch construction algorithm that cycles through data partitions while maintaining per-partition sampling state.

RPR addresses the limitation that empirical risk minimization tends to be dominated by frequent categories. We aim to minimize the ideal loss where each category *c* contributes equally:

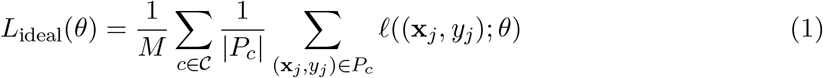

where *P*_*c*_ are data partitions defined by categorical feature combinations and *M* = |𝒞| is the number of partitions.

RPR ensures uniform representation of rare canonical classes while maintaining computational efficiency compared to stratified oversampling approaches [8, 22].

##### Algorithm 1

Round-Robin Partition Rebalancing (RPR)

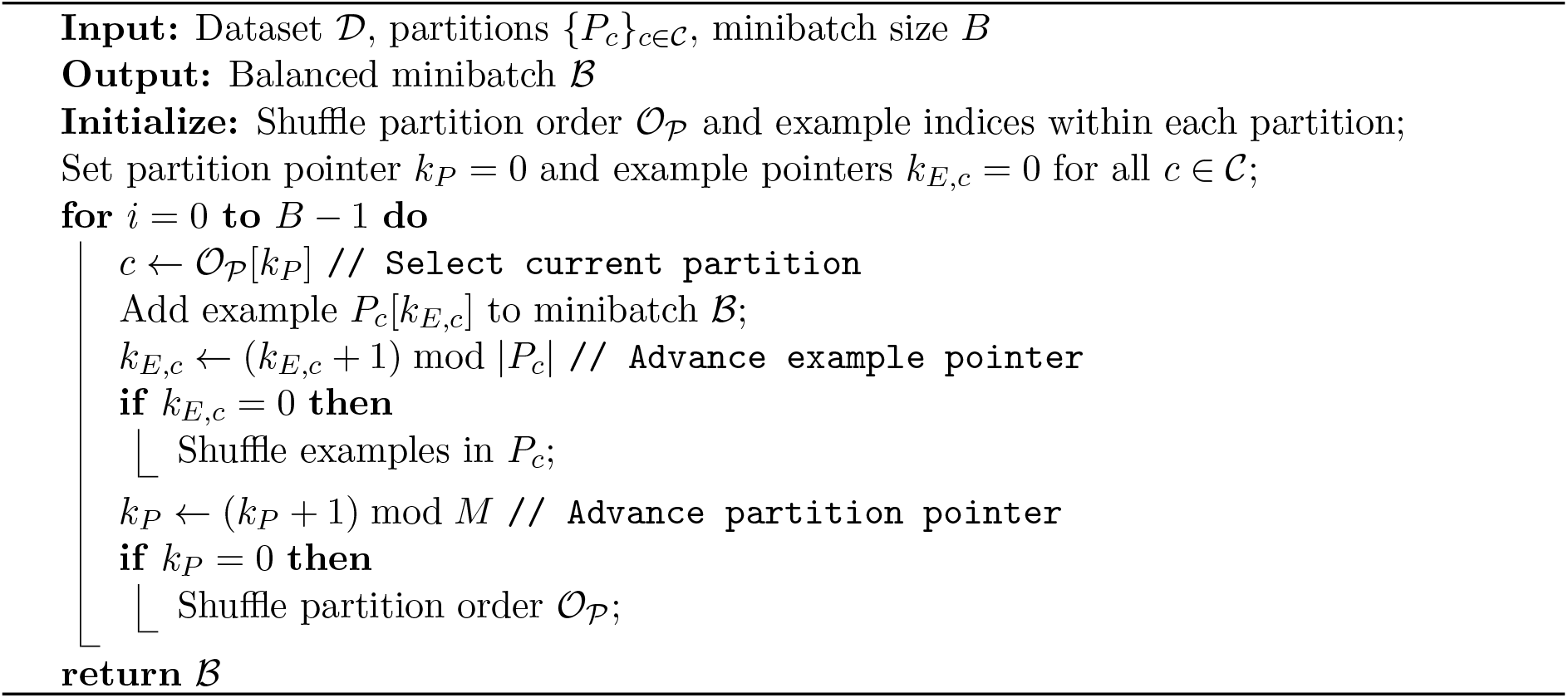

**Multi-Task Training with Label Smoothing** ConformAb employs multi-task learning where each gradient step randomly selects either the diffusion head or one of the CDR classification task heads for optimization. After sampling a task we draw a balanced minibatch from the corresponding task dataset (since the training data for the diffusion head may include unlabeled examples), compute the task loss, and perform a gradient update on the weights of the active subnetwork for the current task. For CDR classification tasks under input corruption, we apply label smoothing with corruption-dependent coefficient:

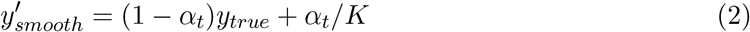

where *α*_*t*_ is the masking fraction, *K* is the number of canonical classes, and *y*_*true*_ is the one-hot true label.

#### 4.3.4 Structure-Guided Generation

ConformAb generates designs that preserve CDR canonical conformations through guided diffusion with explicit steering. We reparameterize the discrete sequence generation problem as a search over continuous hidden states ***h*** that define categorical distributions over tokens when transformed through the final linear token classification head and softmax. Let *H* denote the random variable representing hidden states from the shared encoder *g*(·) that maps sequences to hidden representations. In Algorithm 2 we restate the NOS algorithm for ease of reference. Our approach uses the NOS (diffusio**N O**ptimized **S**ampling) algorithm [21], with the key novelty being our specific choice of value function for CDR canonical conformation preservation. The guidance signal ensures canonical class distributions of the design match those of the seed:

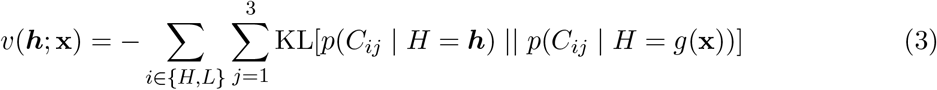

where *p*(*C*_*ij*_ | *H* = ***h***) and *p*(*C*_*ij*_ | *H* = *g*(**x**)) are the predicted canonical class distributions for CDR *j* of chain *i* computed from the guided hidden states and seed hidden states, respectively.

##### Algorithm 2

ConformAb Generation with Diffusio**N O**ptimized **S**ampling (NOS)

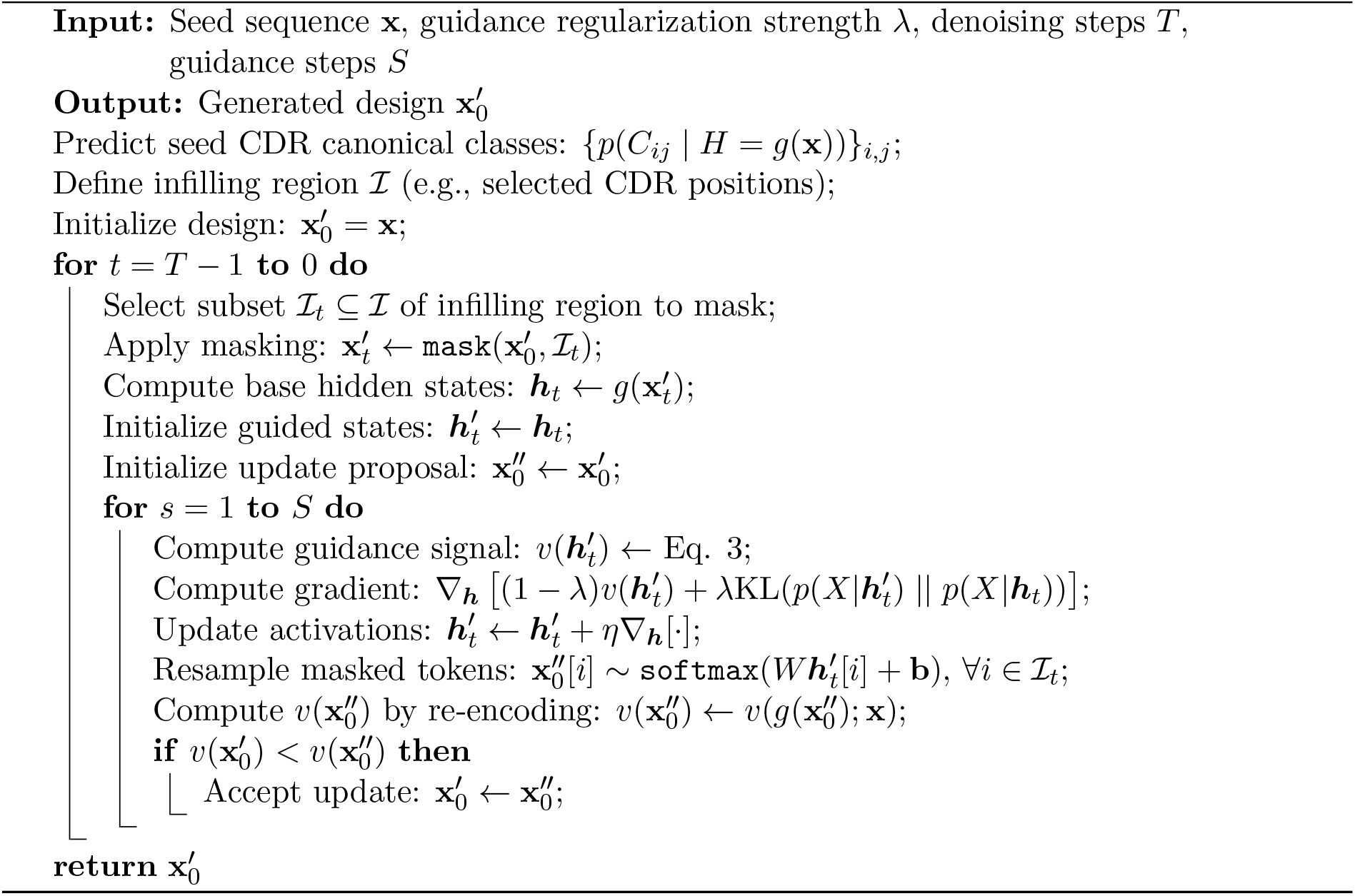

For position selection under edit budget constraints, we use occlusion-based saliency to identify positions most impactful for canonical structure preservation [46].

#### 4.3.5 Implementation Details

Models are implemented using the cortex framework^1^ for modular deep learning in PyTorch. For RPR on each CDR prediction task, we partition data by CDR canonical class. We set the number of partial deep ensemble components for CDR classification to 10. We set the convolution kernel width to 9 in all layers. We set the dimension of the token embeddings to 32, and the dimension of the internal model activations to 256. The shared convolutional encoder was composed of 4 residual CNN blocks, and each of the branch encoders were composed of two residual blocks. We used LayerNorm to normalize the activations and stabilize training.

Our training procedure follows the multi-task partial deep ensemble approach described in [21]. At each gradient step we sample a task head, then sample a minibatch using RPR and compute gradients on the corresponding subnetwork. Since task datasets have different sizes, we parameterize training in terms of total gradient updates rather than epochs.

### 4.4 Antibody antigen binding kinetics measurements

Antibody-antigen binding kinetics were measured by surface plasmon resonance (SPR) on a Biacore 8K instrument (Cytiva) using a multi-cycle kinetics protocol as described in [18]. A standard SPR multi-cycle kinetics experiment was set up using 10 mM HEPES, 150 mM NaCl, 2 mM EDTA, pH 7.4 as running buffer, sensor chip ProA SPR chips for antibody capture, and 10 mM hydrochloric acid pH 1.5 for chip regeneration. Sensorgrams at five different antigen concentrations, ranging from 0 to 100 nM, were collected for each antibody. A typical cycle was defined as follows: 30 seconds of antibody capture using 1 ug/mL antibody solution in SPR running buffer, 180 seconds antigen association, 360 seconds dissociation, and 30 seconds of regeneration. Fit binding kinetic parameters were obtained by fitting a 1:1 binding model to the reference-subtracted data using the Biacore Insight Evaluation software. Binding affinity was defined as the log-transformed equilibrium dissociation constant: 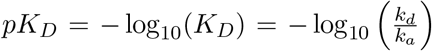, with *k*_*d*_ and *k*_*a*_ representing the dissociation and association rate constants, respectively.

### 4.5 Protein expression, purification, crystallization, and structure determination

Fab constructs for the anti-IL6 and anti-EGFR leads and their design variant were transfected into CHO cells using a 1:2 (HC:LC) DNA ratio, and protein expression was carried out over a period of 10 days. Following expression, the supernatant was collected for purification using GammaBind Plus Sepharose (Cytiva), followed by either size-exclusion chromatography or SP cation exchange chromatography. The resulting purified Fabs were formulated in 20 mM histidine acetate buffer (pH 5.5, 150 mM NaCl). Proteins were concentrated to 10–20 mg/mL, and crystallized [0.2M Ammonium sulfate, 0.1M sodium acetate pH 4.6, 30%w/vPEG2000MME] for the aIL6 lead. Crystals were cryoprotected by addition of 5–15% ethylene glycol. Crystallographic data were collected at the Shanghai Synchrotron Radiation Facility (SSRF) on beamline BL02U1. Data processing and scaling were performed using XDS39, and structures were determined via molecular replacement with PHASER40. The model was built in COOT41, followed by refinement using PHENIX42, with final statistics provided in Supplementary Table S1. Simulated annealing omit difference electron density maps (Supplementary Fig. S1) show the agreement between the final models with the electron density.

### 4.6 Figures

All figures were created using BioRender.com. Antibody structures were visualized using Pymol.

## Acknowledgements

We acknowledge the insightful discussions provided by colleagues at Prescient Design and the Antibody Engineering department at Genentech. We thank Tamica D’Souza for wet lab support and Jia Wu and Kellen Schneider for antibody discovery and immunization experiments. We thank James R. Kiefer, Yanshu Dou and Viva Biotech for structural work and the Shanghai Synchrotron Radiation Facility (SSRF) and staff of beamline BL02U1 for crystallographic data collection. We thank the Genentech NGS core lab for the repertoire sequence data used in this study. We also thank the Prescient Design engineering team and Genentech IT for computing support.

## Author Contribution Statement

I.S., S.S., S.N., and S.K. conceived the work; I.S., S.S., S.A.R., K.Z., J.K., H.M., A.M.W., H.D., R.B., K.C., F.S., V.G., S.K., designed computational experiments. Y.W., S.L., M.D., Y.C., J.B., designed and conducted wet-lab experiments. I.S., S.A.R., S.S., S.K., S.L., analyzed the results. I.S., S.K., S.S., S.A.R., S.L., wrote the initial draft and all authors reviewed the manuscript.

## Additional Information

### Code Availability

ConformAb models are implemented using the cortex framework, available at https://github.com/prescient-design/cortex. Crystallographic data for EGFR and IL6 binders shown in Figure 4 will be deposited in the Protein Data Bank (PDB) and made available upon publication.

## Footnotes

1 https://github.com/prescient-design/cortex

## References

[1] Brennan Abanades, Guy Georges, Alexander Bujotzek, and Charlotte M Deane. Ablooper: fast accurate antibody cdr loop structure prediction with accuracy estimation. Bioinformatics, 38(7): 1877–1880, 2022.

[2] Brennan Abanades, Wing Ki Wong, Fergus Boyles, Guy Georges, Alexander Bujotzek, and Charlotte M Deane. Immunebuilder: Deep-learning models for predicting the structures of immune proteins. Communications Biology, 6(1): 575, 2023.

[3] Jared Adolf-Bryfogle, Oleks Kalyuzhniy, Michael Kubitz, Brian D Weitzner, Xiaozhen Hu, Yumiko Adachi, William R Schief, and Roland L Dunbrack Jr. Rosettaantibodydesign (rabd): A general framework for computational antibody design. PLoS computational biology, 14(4):e1006112, 2018.

[4] Jared Adolf-Bryfogle, Qifang Xu, Benjamin North, Andreas Lehmann, and Roland L Dunbrack Jr. Pyigclassify: a database of antibody cdr structural classifications. Nucleic acids research, 43(D1):D432–D438, 2015.

[5] Bissan Al-Lazikani, Arthur M Lesk, and Cyrus Chothia. Standard conformations for the canonical structures of immunoglobulins. Journal of molecular biology, 273(4): 927–948, 1997.

[6] Jacob Austin, Daniel D Johnson, Jonathan Ho, Daniel Tarlow, and Rianne Van Den Berg. Structured denoising diffusion models in discrete state-spaces. Advances in neural information processing systems, 34: 17981–17993, 2021.

[7] Paul J Carter and Arvind Rajpal. Designing antibodies as therapeutics. Cell, 185(15):2789–2805, 2022.

[8] Nitesh V Chawla, Kevin W Bowyer, Lawrence O Hall, and W Philip Kegelmeyer. Smote: synthetic minority oversampling technique. Journal of artificial intelligence research, 16: 321–357, 2002.

[9] Cyrus Chothia and Arthur M Lesk. Canonical structures for the hypervariable regions of immunoglobulins. Journal of molecular biology, 196(4): 901–917, 1987.

[10] Cyrus Chothia, Arthur M Lesk, Anna Tramontano, Michael Levitt, Sandra J Smith-Gill, Gillian Air, Steven Sheriff, Eduardo A Padlan, David Davies, William R Tulip, et al. Conformations of immunoglobulin hypervariable regions. Nature, 342(6252): 877–883, 1989.

[11] Alexander E Chu, Jinho Kim, Lucy Cheng, Gina El Nesr, Minkai Xu, Richard W Shuai, and Po-Ssu Huang. An all-atom protein generative model. Proceedings of the National Academy of Sciences, 121(27):e2311500121, 2024.

[12] Loredana Lo Conte, Cyrus Chothia, and Joel Janin. The atomic structure of protein-protein recognition sites. Journal of molecular biology, 285(5): 2177–2198, 1999.

[13] Justas Dauparas, Ivan Anishchenko, Nathaniel Bennett, Hua Bai, Robert J Ragotte, Lukas F Milles, Basile IM Wicky, Alexis Courbet, Rob J de Haas, Neville Bethel, et al. Robust deep learning–based protein sequence design using proteinmpnn. Science, 378(6615): 49–56, 2022.

[14] James Dunbar, Konrad Krawczyk, Jinwoo Leem, Terry Baker, Angelika Fuchs, Guy Georges, Jiye Shi, and Charlotte M Deane. Sabdab: the structural antibody database. Nucleic acids research, 42(D1):D1140–D1146, 2014.

[15] Monica L Fernandez-Quintero, Katharina B Kroell, Florian Hofer, Jakob R Riccabona, and Klaus R Liedl. Mutation of framework residue h71 results in different antibody paratope states in solution. Frontiers in Immunology, 12:630034, 2021.

[16] Noelia Ferruz, Steffen Schmidt, and Birte Hocker. Protgpt2 is a deep unsupervised language model for protein design. Nature communications, 13(1): 4348, 2022.

[17] Nathan C Frey, Dan Berenberg, Joseph Kleinhenz, Isidro Hotzel, Julien Lafrance-Vanasse, Ryan Lewis Kelly, Yan Wu, Arvind Rajpal, Stephen Ra, Richard Bonneau, et al. Learning protein family manifolds with smoothed energy-based models. In ICLR 2023 Workshop on Physics for Machine Learning, 2023.

[18] Nathan C Frey, Isidro Hotzel, Samuel D Stanton, Ryan L Kelly, Robert G Alberstein, Emily K Makowski, Karolis Martinkus, Dan Berenberg, Jack Bevers III, Tyler Bryson, et al. Lab-in-the-loop therapeutic antibody design with deep learning. bioRxiv, pages 2025–02, 2025.

[19] Vladimir Gligorijevic, Daniel Berenberg, Stephen Ra, Andrew Watkins, Simon Kelow, Kyunghyun Cho, and Richard Bonneau. Function-guided protein design by deep manifold sampling. bioRxiv, pages 2021–12, 2021.

[20] Alexander Greenshields-Watson, Brennan Abanades, and Charlotte M Deane. Investigating the ability of deep learning-based structure prediction to extrapolate and/or enrich the set of antibody cdr canonical forms. Frontiers in Immunology, 15:1352703, 2024.

[21] Nate Gruver, Samuel Stanton, Nathan C Frey, Tim GJ Rudner, Isidro Hotzel, Julien Lafrance-Vanasse, Arvind Rajpal, Kyunghyun Cho, and Andrew Gordon Wilson. Protein design with guided discrete diffusion. arXiv preprint arXiv:2305.20009, 2023.

[22] Haibo He and Edwardo A Garcia. Learning from imbalanced data. IEEE Transactions on knowledge and data engineering, 21(9): 1263–1284, 2009.

[23] Emiel Hoogeboom, Didrik Nielsen, Priyank Jaini, Patrick Forre, and Max Welling. Argmax flows and multinomial diffusion: Learning categorical distributions. In Advances in Neural Information Processing Systems, volume 34, pages 12454–12465, 2021.

[24] Mark Hutchinson, Jeffrey A Ruffolo, Nantaporn Haskins, Michael Iannotti, Giuliana Vozza, Tony Pham, Nurjahan Mehzabeen, Harini Shandilya, Keith Rickert, Rebecca Croasdale-Wood, et al. Toward enhancement of antibody thermostability and affinity by computational design in the absence of antigen. In MAbs, volume 16, page 2362775. Taylor & Francis, 2024.

[25] John Ingraham, Vikas Garg, Regina Barzilay, and Tommi Jaakkola. Generative models for graph-based protein design. Advances in neural information processing systems, 32, 2019.

[26] Kaiyi Jiang, Zhaoqing Yan, Matteo Di Bernardo, Samantha R Sgrizzi, Lukas Villiger, Alisan Kayabolen, Byungji Kim, Josephine K Carscadden, Masahiro Hiraizumi, Hiroshi Nishimasu, et al. Rapid protein evolution by few-shot learning with a protein language model. bioRxiv, 2024.

[27] Wengong Jin, Jeremy Wohlwend, Regina Barzilay, and Tommi Jaakkola. Iterative refinement graph neural network for antibody sequence-structure co-design. arXiv preprint arXiv:2110.04624, 2021.

[28] Simon Kelow, Bulat Faezov, Qifang Xu, Mitchell Parker, Jared Adolf-Bryfogle, and Roland L Dunbrack Jr. A penultimate classification of canonical antibody cdr conformations. bioRxiv, pages 2022–10, 2022.

[29] Aleksandr Kovaltsuk, Jinwoo Leem, Sebastian Kelm, James Snowden, Charlotte M Deane, and Konrad Krawczyk. Observed antibody space: a resource for data mining nextgeneration sequencing of antibody repertoires. The Journal of Immunology, 201(8):2502–2509, 2018.

[30] Zeming Lin, Halil Akin, Roshan Rao, Brian Hie, Zhongkai Zhu, Wenting Lu, Allan dos Santos Costa, Maryam Fazel-Zarandi, Tom Sercu, Sal Candido, et al. Language models of protein sequences at the scale of evolution enable accurate structure prediction. BioRxiv, 2022:500902, 2022.

[31] Sidney Lyayuga Lisanza, Jacob Merle Gershon, Sam Wayne Kenmore Tipps, Lucas Arnoldt, Samuel Hendel, Jeremiah Nelson Sims, Xinting Li, and David Baker. Joint generation of protein sequence and structure with rosettafold sequence space diffusion. bioRxiv, pages 2023–05, 2023.

[32] Ruei-Min Lu, Yu-Chyi Hwang, I-Ju Liu, Chi-Chiu Lee, Han-Zen Tsai, Hsin-Jung Li, and Han-Chung Wu. Development of therapeutic antibodies for the treatment of diseases. Journal of biomedical science, 27: 1–30, 2020.

[33] Ali Madani, Bryan McCann, Nikhil Naik, Nitish Shirish Keskar, Namrata Anand, Raphael R Eguchi, Po-Ssu Huang, and Richard Socher. Progen: Language modeling for protein generation. arXiv preprint arXiv:2004.03497, 2020.

[34] Sai Pooja Mahajan, Jeffrey A Ruffolo, Rahel Frick, and Jeffrey J Gray. Hallucinating structure-conditioned antibody libraries for target-specific binders. Frontiers in immunology, 13:999034, 2022.

[35] Andrew CR Martin and Janet M Thornton. Structural families in loops of homologous proteins: automatic classification, modelling and application to antibodies. Journal of molecular biology, 263(5): 800–815, 1996.

[36] Dimitris Nikoloudis, Jim E Pitts, and Jose W Saldanha. A complete, multi-level conformational clustering of antibody complementarity-determining regions. PeerJ, 2:e456, 2014.

[37] Benjamin North, Andreas Lehmann, and Roland L Dunbrack Jr. A new clustering of antibody cdr loop conformations. Journal of molecular biology, 406(2): 228–256, 2011.

[38] Jaroslaw Nowak, Terry Baker, Guy Georges, Sebastian Kelm, Stefan Klostermann, Jiye Shi, Sudharsan Sridharan, and Charlotte M Deane. Length-independent structural similarities enrich the antibody cdr canonical class model. In MAbs, volume 8, pages 751–760. Taylor & Francis, 2016.

[39] Baldomero Oliva, Paul A Bates, Enrique Querol, Francesc X Aviles, and Michael JE Sternberg. Automated classification of antibody complementarity determining region 3 of the heavy chain (h3) loops into canonical forms and its application to protein structure prediction. Journal of molecular biology, 279(5): 1193–1210, 1998.

[40] Tobias H Olsen, Fergus Boyles, and Charlotte M Deane. Observed antibody space: A diverse database of cleaned, annotated, and translated unpaired and paired antibody sequences. Protein Science, 31(1): 141–146, 2022.

[41] Sergey Ovchinnikov and Po-Ssu Huang. Structure-based protein design with deep learning. Current opinion in chemical biology, 65: 136–144, 2021.

[42] Alexander Rives, Joshua Meier, Tom Sercu, Siddharth Goyal, Zeming Lin, Jason Liu, Demi Guo, Myle Ott, C Lawrence Zitnick, Jerry Ma, et al. Biological structure and function emerge from scaling unsupervised learning to 250 million protein sequences. Proceedings of the National Academy of Sciences, 118(15):e2016239118, 2021.

[43] Hiroki Shirai, Akinori Kidera, and Haruki Nakamura. Structural classification of cdr-h3 in antibodies. FEBS letters, 399(1–2): 1–8, 1996.

[44] Yang Song, Prafulla Dhariwal, Mark Chen, and Ilya Sutskever. Consistency models. In International Conference on Machine Learning, pages 32211–32252. PMLR, 2023.

[45] Anna Tramontano, Cyrus Chothia, and Arthur M Lesk. Framework residue 71 is a major determinant of the position and conformation of the second hypervariable region in the vh domains of immunoglobulins. Journal of molecular biology, 215(1): 175–182, 1990.

[46] Amy Wang, Zhe Sang, Samuel D. Stanton, Jennifer L. Hofmann, Saeed Izadi, Eliott Park, Jan Ludwiczak, Matthieu Kirchmeyer, Darcy Davidson, Andrew Maier, Tom Pritsky, Nathan C. Frey, Andrew M. Watkins, and Franziska Seeger. A guided design framework for the optimization of therapeutic-like antibodies. In Proceedings of the GEM Workshop at the International Conference on Learning Representations (ICLR), 2025. GEM Workshop.

[47] Joseph L Watson, David Juergens, Nathaniel R Bennett, Brian L Trippe, Jason Yim, Helen E Eisenach, Woody Ahern, Andrew J Borst, Robert J Ragotte, Lukas F Milles, et al. De novo design of protein structure and function with rfdiffusion. Nature, pages 1–3, 2023.

[48] George J Weiner. Building better monoclonal antibody-based therapeutics. Nature Reviews Cancer, 15(6): 361–370, 2015.

[49] Nicholas Whitelegg and Anthony R Rees. Antibody variable regions: toward a unified modeling method. Antibody engineering: methods and protocols, pages 51–91, 2004.

[50] Wing Ki Wong, Guy Georges, Francesca Ros, Sebastian Kelm, Alan P Lewis, Bruck Taddese, Jinwoo Leem, and Charlotte M Deane. Scalop: sequence-based antibody canonical loop structure annotation. Bioinformatics, 35(10): 1774–1776, 2019.

[51] Yilun Xu, Mingyang Liu, Niloofar Tehrani, Yaodong Duan, and Tommi Jaakkola. Restart sampling for improving generative processes. In Advances in Neural Information Processing Systems, 2023.

